# Enterococcus faecalis modulates phase variation in Clostridioides difficile

**DOI:** 10.1101/2025.07.24.666506

**Authors:** Ashley S. Weiss, Jilarie A. Santos-Santiago, Orlaith Keenan, Alexander B. Smith, Montana Knight, Joseph P. Zackular, Rita Tamayo

**Affiliations:** Division of Protective Immunity, Children’s Hospital of Philadelphia, Philadelphia, PA, USA; Department of Microbiology and Immunology, University of North Carolina Chapel Hill, Chapel Hill, North Carolina, USA; Department of Biomedical and Health Informatics, Children’s Hospital of Philadelphia, Philadelphia, Pennsylvania, USA; Department of Pathology and Laboratory Medicine, Perelman School of Medicine, University of Pennsylvania; Philadelphia, PA, USA; Center for Microbial Medicine, Children’s Hospital of Philadelphia, Philadelphia, PA, USA

## Abstract

To adapt and persist in the gastrointestinal tract, many enteric pathogens, including *Clostridioides difficile,* employ strategies such as phase variation to generate phenotypically heterogeneous populations. Notably, the role of the gut microbiota and polymicrobial interactions in shaping population heterogeneity of invading pathogens has not been explored. Here, we show that *Enterococcus faecalis*, an opportunistic pathogen that thrives in the inflamed gut during *C. difficile* infection, can impact the phase variable CmrRST signal transduction system in *C. difficile*. The CmrRST system controls multiple phenotypes including colony morphology, cell elongation, and cell chaining in *C. difficile*. Here we describe how interactions between *E. faecalis* and *C. difficile* on solid media lead to a marked shift in *C. difficile* phenotypes associated with phase variation of CmrRST. Specifically, *E. faecalis* drives a switch of the *C. difficile* population to the *cmr-*ON state leading to chaining and a rough colony morphology. This phenomenon preferentially occurs with *E. faecalis* among the enterococci, as other enterococcal species do not show a similar effect, suggesting that the composition of the polymicrobial environment in the gut is likely critical to shaping *C. difficile* population heterogeneity. Our findings shed light on the complex role that microbial ecology and polymicrobial interactions can have in the phenotypic heterogeneity of invading pathogens.

**IMPORTANCE STATEMENT:** *Clostridioides difficile* is an enteric pathogen with critical implications for public health. The microbial ecosystem in which *C. difficile* resides shapes the behavior and fitness of *C. difficile*; however, the mechanisms underlying these interactions are not well defined. Here, we demonstrate that *Enterococcus faecalis*, an opportunistic pathogen known to co-colonize the gut with *C. difficile*, influences phase variation and downstream growth phenotypes in *C. difficile*. This phenomenon represents a new paradigm by which co-residing bacteria can modulate phase variation dynamics in *C. difficile* or other enteric pathogens. Understanding factors that influence *C. difficile* behavior may elucidate new therapeutic strategies, especially in complex polymicrobial infections.

## INTRODUCTION

*Clostridioides difficile* is a major nosocomial bacterial pathogen that causes hundreds of thousands of healthcare and community-acquired infections each year in the United States (1). *C. difficile* infection (CDI) ranges from mild diarrhea to potentially fatal toxic megacolon (1–3). Perturbations of the gut microbiota, usually through antibiotic use, diminish bacterial-driven colonization resistance and facilitate colonization by *C. difficile* in the gastrointestinal (GI) tract, leading to CDI (3). The GI tract of humans and other mammals harbors a dynamic and diverse microbial ecosystem populated by a rich collection of microorganisms; however, the mechanisms by which the gut microbiota shape *C. difficile* fitness and virulence remain understudied. Bacterial phase variation allows microbial populations to adapt to sudden environmental fluctuations more rapidly than conventional gene mutations (4, 5). Phase variation is a bet-hedging process that allows for stochastic shifting of phenotypes, generating phenotypically heterogeneous populations that are better equipped to survive abrupt stresses. Many *C. difficile* strains exhibit phase variation of colony morphology, shifting between rough and smooth colony morphotypes (6, 7). The rough and smooth morphotypes differ in additional phenotypes, including cell length, chaining, swimming, surface motility, biofilm production, and virulence in a hamster model of CDI (7).

Alternation between rough and smooth colony morphotypes is mediated by the reversible inversion of the “*cmr* switch”, a regulatory DNA sequence upstream of the <u>c</u>olony <u>m</u>orphology <u>r</u>egulators RST (*cmrRST)* operon encoding a signal transduction system (7–9). The *cmr* switch contains a promoter and undergoes inversion via the activity of the site-specific tyrosine DNA recombinase RecV (8, 10). One orientation of the *cmr* switch promotes *cmrRST* transcription and yields the rough colony morphotype (*cmr*-ON state). The inverse orientation does not stimulate expression, leading to the smooth colony morphotype (*cmr*-OFF state). Alongside the promoter within the *cmr* switch, *cmrRST* is regulated by two additional promoters: the furthest upstream promoter leads to a transcript containing a cyclic diguanylate (c-di-GMP) riboswitch, and one is positively autoregulated by CmrR (7, 9). Under basal c-di-GMP conditions, *cmr* switch-mediated modulation of *cmrRST* expression is sufficient to phase vary colony morphology (9). Importantly, the environmental conditions that influence the phase variation of colony morphology are unknown.

Polymicrobial interactions are critical to the pathogenesis of numerous infections. One of the most dominant members of the gut microbiota of patients with CDI is enterococci (11, 12). Vancomycin-resistant *Enterococcus* (VRE) commonly co-infects patients with *C. difficile*, and enterococci are enriched in the *C. difficile*-infected gut (11, 12). Previously, we reported that enterococci enhance *C. difficile* colonization and pathogenesis during infection through metabolic interactions in the gut (11). However, the molecular mechanisms by which enterococci alter *C. difficile* behavior have not been comprehensively defined.

In this study, we evaluated the effect of *E. faecalis* OG1RF on *C. difficile* R20291 phenotypes associated with the *cmr* switch. We show that *E. faecalis* enriches the *cmr-* ON state in the *C. difficile* population. We also show that this enrichment is preferential to *E. faecalis* strains compared to other enterococci. Together, this work establishes a new paradigm in which the gut microbiota shapes the behavior of invading pathogens through phase variation.

## RESULTS

### *E. faecalis* OG1RF stimulates *C. difficile* R20291 surface motility independently from flagella and type IV pili

*C. difficile* is exposed to a dynamic, polymicrobial ecosystem in the gut during infection. In our previous work, we demonstrated that a group of opportunistic pathogens, the enterococci, markedly impacts the outcome of CDI. Specifically, we found that *E. faecalis* increases *C. difficile* fitness and virulence through metabolic crosstalk (11). Notably, we also observed that these two organisms interacted closely at the mucosal surface during infection, suggesting that polymicrobial interactions between these pathogens on surfaces may be critical. To explore this in greater depth, we grew mixed species cultures on agar surfaces with *C. difficile* strain R20291 and *E. faecalis* strain OG1RF (**Figure 1A**). We noted a dramatic shift in macrocolony morphology and the formation of tendril-like structures in the *C. difficile* outer ring of the colony. *C. difficile* and *E. faecalis* localization was determined by selective plating of the inner and outer portions of the macrocolonies. To evaluate if this shift in morphology was contact-dependent, we next performed a proximity assay with *C. difficile* and *E. faecalis* at gradually increasing distance (**Figure 1B**). We observed that *E. faecalis* stimulates changes in *C. difficile* when the two bacteria are plated in close proximity, with tendrils stretching from *C. difficile* towards *E. faecalis* in a directional manner. This effect between *C. difficile* and *E. faecalis* was enhanced with greater proximity. These results indicate that *E. faecalis* may consume or secrete a soluble factor that alters growth phenotypes in *C. difficile*.

**Figure 1.**
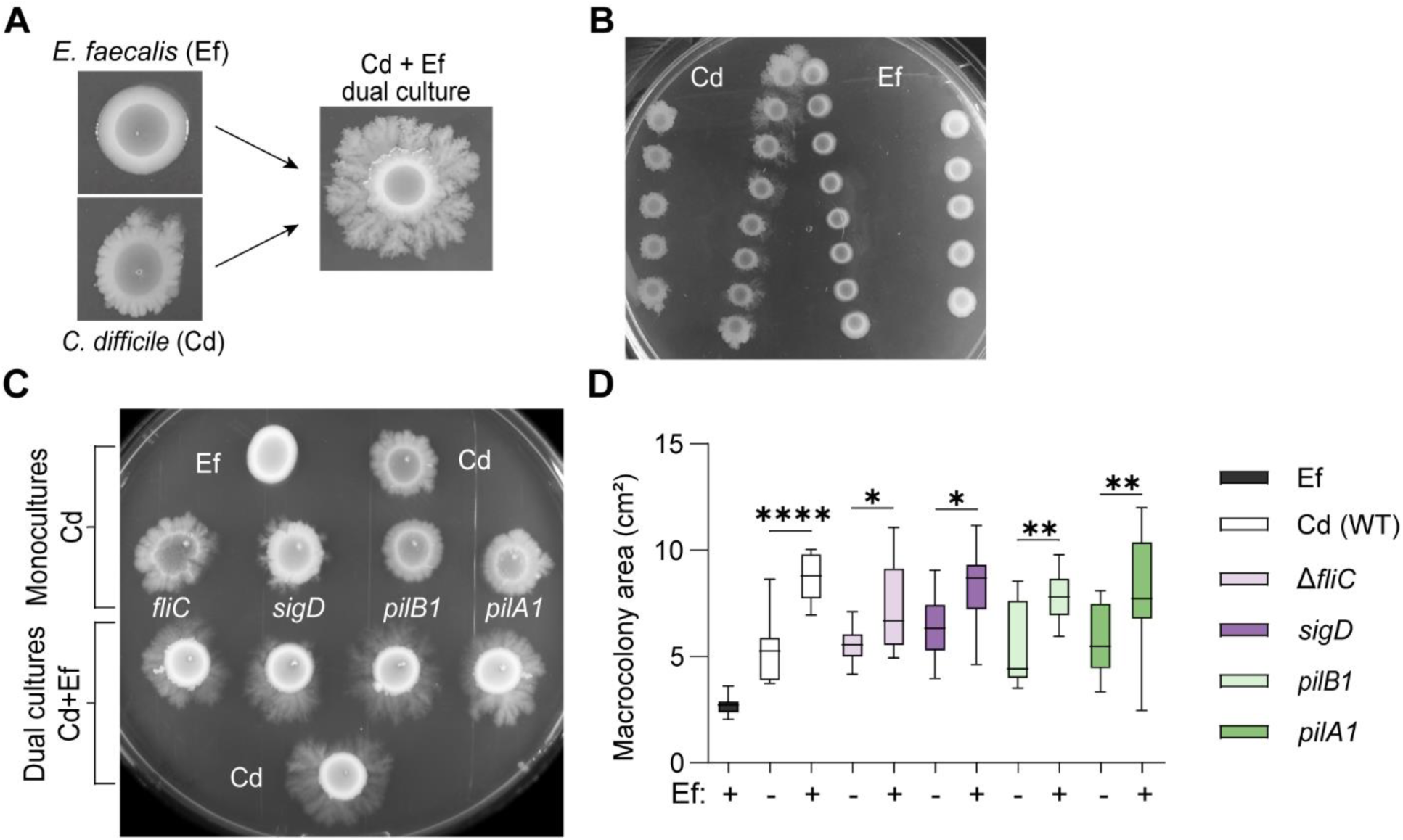
*E. faecalis* stimulates surface motility in *C. difficile* independently of flagella and type IV pili. (**A**) Representative images of macrocolonies after 7 days of growth. Wildtype (WT) *C. difficile* (Cd), *E. faecalis* (Ef), and dual culture with Cd and Ef at a 1:1 ratio. (**B**) Proximity growth assay. Representative image of Cd and Ef monocultures after 7 days of growth on BHIS agar. (**C**) Representative image of surface motility after 7 days of growth. Surface motility of Ef, WT Cd, Δ*fliC* Cd, *sigD::ermB* Cd, *pilB1::ermB* Cd, and *pilA1::ermB* Cd in monocultures (top) or dual cultures (bottom) with Ef at a 1:1 ratio. (**D**) Macrocolony area on day 7 of surface growth. Data from independent experiments is presented as means and standard error. **** p < 0.0001, ** p < 0.01, * p < 0.05, one-way ANOVA with Šídák’s multiple comparison test.

In many bacterial species, tendril formation and surface motility are mediated by pili and flagella (13–15). To determine the potential contribution of pili and flagellar structures in surface motility of *C. difficile* in response to *E. faecalis*, *C. difficile* mutants lacking type IV pili (T4P) (*pilA1::ermB*, *pilB1::ermB*) or flagella (Δ*fliC, sigD::ermB*) were used in motility assays (**Figure 1C-D**) (16, 17). Monocultures of wildtype and mutant *C. difficile* without *E. faecalis* were included as controls. All *C. difficile* strains tested displayed increased surface motility in dual culture with *E. faecalis* (bottom) relative to *C. difficile* monocultures (top). Quantification of macrocolony growth area showed significantly greater surface motility for *C. difficile* wildtype and mutant strains in co-culture with *E. faecalis* after 7 days (**Figure 1D**), indicating that the observed increase in surface motility is independent of flagella and T4P. Since *E. faecalis* is a nonmotile bacterium, the increase in macrocolony area in dual cultures is only attributable to *C. difficile* motility changes, not to *E. faecalis* movement or growth (18).

### The *E. faecalis* OG1RF-mediated increase in surface motility of *C. difficile* R20291 requires *cmrT*

We previously determined that the signal transduction system CmrRST promotes *C. difficile* surface motility in a flagellum- and T4P-independent manner (7). We also demonstrated that CmrR and CmrS are dispensable for CmrRST-dependent surface migration in *C. difficile*, while CmrT is required (7, 9). Thus, we asked if the CmrRST system is involved in the phenotypic shifts in *C. difficile* when exposed to *E. faecalis* on solid surfaces. First, we evaluated the impact of *E. faecalis* on the transcription of *cmrRST* in wildtype *C. difficile*. After growth in monoculture or dual culture for 5 days, we measured *cmrR*, *cmrS*, and *cmrT* transcript abundance using qRT-PCR (**Figure 2A**). We observed a significantly higher transcript abundance of all three genes in co-culture with *E. faecalis* compared to the *C. difficile* monoculture.

**Figure 2.**
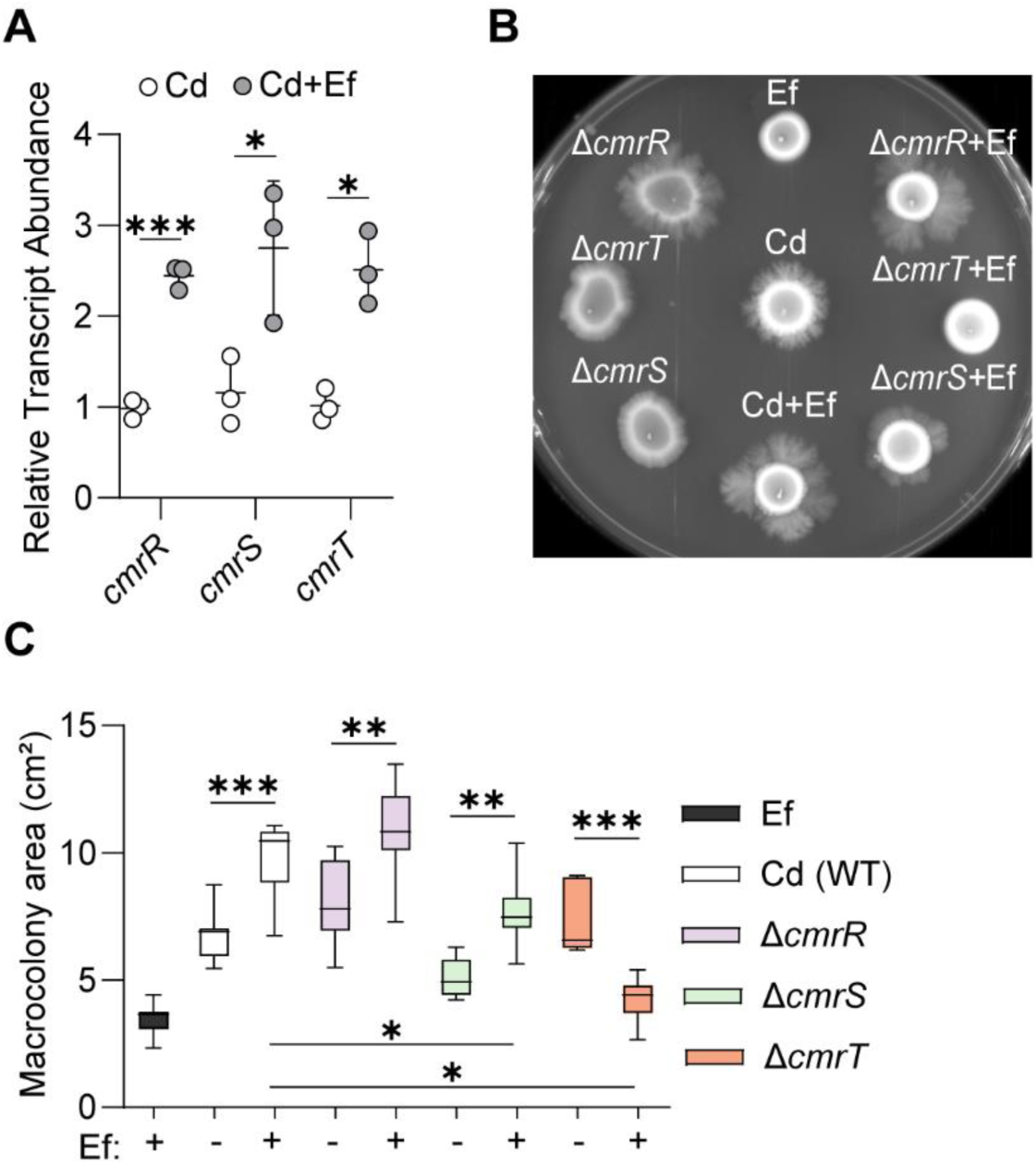
*C. difficile* requires *cmrT* for the *E. faecalis*-mediated increase in surface motility. (**A**) Transcription of *cmrRST* in *C. difficile*. Transcript abundance of *cmrR*, *cmrS*, and *cmrT* from *C. difficile* in monoculture and dual culture after 5 days of growth on BHIS agar. (**B**) Representative image of surface motility after 7 days of growth on BHIS agar. Surface motility of Ef, Cd, Δ*cmrR* Cd, Δ*cmrS* Cd, and Δ*cmrT* Cd in monocultures or dual cultures with Ef at a 1:1 ratio. (**C**) Macrocolony area on day 7 of surface growth. Data is presented as means and standard error, with symbols representing individual values from independent experiments. *** p < 0.001, ** p < 0.01, p < 0.05, Welch’s t test (A) and one-way ANOVA with Šídák’s multiple comparison test (C).

Next, we performed surface motility assays and quantified the macrocolony area with *C. difficile* strains lacking the individual genes of *cmrRST*, including each of the response regulator genes (Δ*cmrR*, Δ*cmrT*) and the histidine kinase gene (Δ*cmrS*) (**Figure 2B-C**). After 7 days on agar in dual culture with *E. faecalis,* the *C. difficile* Δ*cmrR* and Δ*cmrS* mutants showed increased surface motility compared to monoculture controls, indicating that *cmrR* and *cmrS* are not required for the effect of *E. faecalis* on the surface migration of *C. difficile* (**Figure 2B**). In contrast, the Δ*cmrT* mutant displayed decreased surface motility with *E. faecalis*. The lack of surface spreading in Δ*cmrT* was not due to *C. difficile* death, as the Δ*cmrT* mutant was determined to be viable in monoculture and dual culture at day 7 by plating (data not shown). These observations are consistent with our previous work showing that CmrT is required for CmrRST-mediated surface motility (7, 9). Together, these results point to the involvement of the CmrRST system and the importance of *cmrT* in the *E. faecalis*-mediated increase in surface motility of *C. difficile*.

### *E. faecalis* OG1RF promotes *C. difficile* R20291 surface motility through the *cmr* switch

We next sought to determine the regulatory mechanism in *C. difficile* by which *E. faecalis* alters *cmrRST* expression and surface motility phenotypes. Regulation of CmrRST is complex, with multiple levels of transcriptional control and likely post-translational control via phosphotransfer between the histidine kinase and response regulator. Expression of the operon is regulated by the *cmr* switch, levels of c-di-GMP, and autoregulatory activity of CmrR (7, 9, 19). We used a panel of *C. difficile* mutants to determine the contribution of each CmrRST regulatory mechanism. First, as noted above, deletion of *cmrR* did not affect the increase in surface motility of *C. difficile* mediated by *E. faecalis*, (**Figure 2B-C**), indicating that the autoregulatory activity of CmrR is dispensable for the phenotype.

We next analyzed the orientation of the *cmr* switch in mono- and dual cultures of *C. difficile* and *E. faecalis* cultured on agar media using qPCR to determine the proportions of *cmr*-ON and *cmr*-OFF *C. difficile* cells in a population (7, 8). We observed a significant increase in the proportion of *cmr-*ON *C. difficile* when in dual culture (85.2%) by day 3 of growth (**Figure 3A**). In contrast, *C. difficile* monoculture remained around 31.6% *cmr-*ON during the same time frame. These results indicate that *E. faecalis* impacts CmrRST phase variation, enriching for *cmr*-ON *C. difficile*.

**Figure 3.**
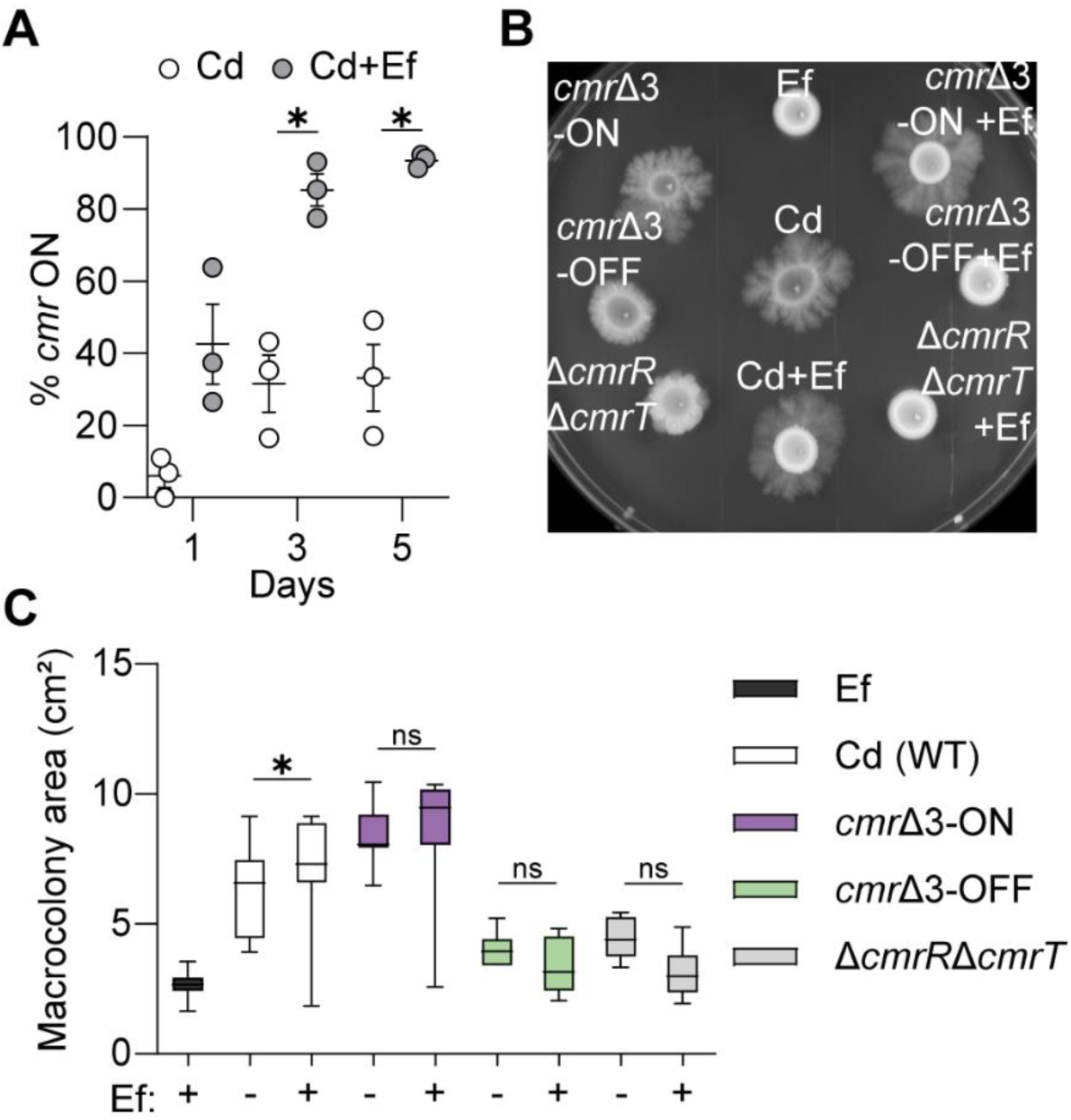
*E. faecalis* promotes surface motility of *C. difficile* through *cmrRST*. (**A**) Quantitative PCR of *cmr* switch orientation in macrocolonies of Cd or Cd and Ef at days 1, 3, and 5 after incubation. Data from 3 biological replicates were analyzed. (**B**) Representative image of surface motility after 5 days of growth. Surface motility of Ef, Cd, *cmr*Δ3-ON Cd, *cmr*Δ3-OFF Cd, and Δ*cmrR*Δ*cmrT* Cd in monocultures or dual cultures with WT Ef at a 1:1 ratio. (**C**) Macrocolony area after 5 days of surface motility growth. Data from independent experiments is presented as means and standard error. * p < 0.05, unpaired t-tests (A) and one-way ANOVA with Šídák’s multiple comparison test (C).

Since *cmrT* is required for increasing *C. difficile* surface motility (Figure 2C), we also investigated whether *cmrT* was required for the *E. faecalis*-mediated increase in *cmr*-ON *C. difficile*. While the *C. difficile* Δ*cmrR* and Δ*cmrS* mutants shifted towards *cmr*-ON in monoculture and dual culture with *E. faecalis*, the Δ*cmrT* mutant did not display an increase in *cmr*-ON, indicating *cmrT* is indispensable for this shift (**Supplemental Figure 1**).

Next, we directly evaluated the role of the *cmr* switch and phase variation of CmrRST in surface motility. We conducted surface motility assays with previously described *C. difficile* mutants that are unable to invert the *cmr* switch and are thus phase-locked in the ON (*cmr*Δ3-ON) or OFF (*cmr*Δ3-OFF) orientation (**Figure 3B-C**) (9). We also included a *C. difficile* mutant lacking both response regulators, Δ*cmrR*Δ*cmrT*, as a control. We predicted that without the ability to invert the *cmr* switch, *E. faecalis* would be unable to stimulate *C. difficile* surface motility. As expected, there were no significant differences in macrocolony area between monocultures and dual cultures of *cmr*Δ3-ON, *cmr*Δ3-OFF, and Δ*cmrR*Δ*cmrT* mutants with *E. faecalis* as early as day 5.

We previously showed that increased c-di-GMP promotes *C. difficile* surface motility independent of CmrRST phase variation, presenting an alternative, non-mutually exclusive mechanism by which *E. faecalis* augments *C. difficile* surface motility (20). However, the absence of differences between *cmr*Δ3-OFF in *C. difficile* monoculture and dual culture with *E. faecalis* suggests that c-di-GMP levels are unlikely to contribute (**Figure 2A**). Nonetheless, we directly evaluated this possibility using *C. difficile* containing a c-di-GMP riboswitch biosensor (p*_gluD-_*PRS::mCherryOpt) (**Supplemental Figure 2**) (20). No differences in fluorescence were observed between monocultures of *C. difficile* and dual cultures with *E. faecalis*, supporting the conclusion that global c-di-GMP level does not play a significant role. Taken together, these results indicate that *E. faecalis*-stimulated surface motility occurs primarily via the *cmr* switch, and not through CmrR autoregulation or shifts in global c-di-GMP levels.

### *E. faecalis* OG1RF enrichment of *cmr*-ON *C. difficile* R20291 promotes *cmr*-ON-associated phenotypes

The orientation of the *cmr*-switch impacts multiple additional phenotypes in *C. difficile,* whereby the *cmr*-ON stain corresponds to greater cell length, cell chaining, and a rough colony morphology (7). Because of the shift towards a *cmr-* ON population in *C. difficile* in dual culture with *E. faecalis*, we predicted that *E. faecalis* influences these additional CmrRST-associated phenotypes. First, we evaluated cell morphology by conducting scanning electron microscopy (SEM) on macrocolonies grown on agar medium for 7 days (**Figure 4A**). We imaged dual cultures of *E. faecalis* with wildtype *C. difficile* or the Δ*cmrR*Δ*cmrT* mutant and observed longer bacilli and chained cells in *C. difficile* wildtype relative to the mutant. We further characterized cell length as well as chaining by Gram staining cells collected from monoculture and dual culture macrocolonies of wildtype or Δ*cmrR*Δ*cmrT C. difficile* with *E. faecalis* (**Figure 4B**). Cell chaining was only apparent for *C. difficile* in dual culture. Quantification of the length of the bacillus from wildtype and Δ*cmrR*Δ*cmrT C. difficile* in monoculture and dual culture with *E. faecalis* revealed significantly greater length of the wildtype *C. difficile*, and not Δ*cmrR*Δ*cmrT,* in dual cultures relative to monocultures but not in (**Figure 4C**). Finally, we visually examined the colony morphology of *C. difficile* in monoculture or dual cultures with *E. faecalis* and observed a higher proportion of rough colonies in dual cultures (**Figure 4D-E**). Together, these data suggest that *E. faecalis* promotes phenotypes associated with the *cmr-*ON state in *C. difficile*.

**Figure 4.**
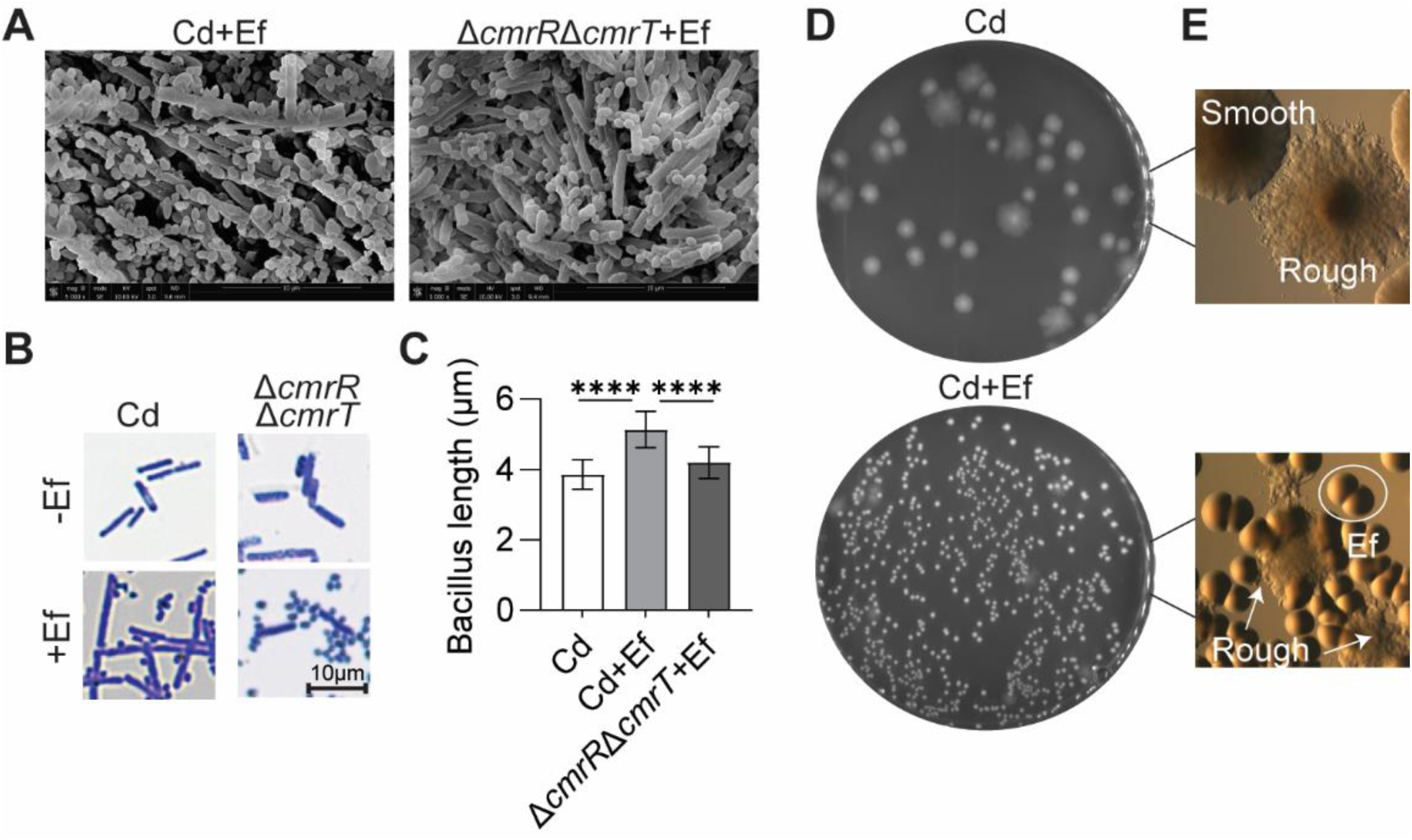
*E. faecalis* stimulates *cmr*-associated phenotypes in *C. difficile*. (**A**) SEM images from dual cultures of Cd or Δ*cmrR*Δ*cmrT* Cd with Ef at a 1:1 ratio after 7 days of growth on surface motility plates. (**B**) Gram stains of Cd and Δ*cmrR*Δ*cmrT* Cd in monocultures and dual cultures with Ef at a 1:1 ratio after 3 days of growth on BHIS plates. Shown are representative images taken at 60X. (**C**) Quantification of cell length of Cd and Δ*cmrR*Δ*cmrT* in mono- or dual-culture culture with Ef at a 1:1 ratio after 3 days of growth on BHIS plates. Data presented as means with SD. **** p < 0.0001, two-way ANOVA. (**D-E**) Representative images of plates (D) and colony morphology (E) of Cd monoculture and dual culture with Ef after 7 days of growth.

### *E. faecalis* strains and specific enteric bacteria increase *cmr-*ON phenotypes in select *C. difficile* strains

CmrRST is highly conserved across *C. difficile* strains of divergent ribotypes (7, 9). Thus, we tested if *E. faecalis* could promote *cmr*-ON phenotypes, specifically increased surface motility, in other *C. difficile* clades and ribotypes aside from R20291 (RT027). We found that UK1 (RT027) and ATCC BAA1875 (RT078) showed a significant increase in surface motility when co-cultured with *E. faecalis* OG1RF (**Figure 5A-B**). The *C. difficile* M120 (RT078) strain did not exhibit a substantial increase in spreading within the same time frame, although we observed an increase in tendril formation. Interestingly, when *C. difficile* strains from other ribotypes, 630 (RT012), VPI10463 (RT087), and ATCC 43598 (RT017), were co-cultured with *E. faecalis* OG1RF, no significant changes in surface motility were observed (**Figure 5C-D**).

**Figure 5.**
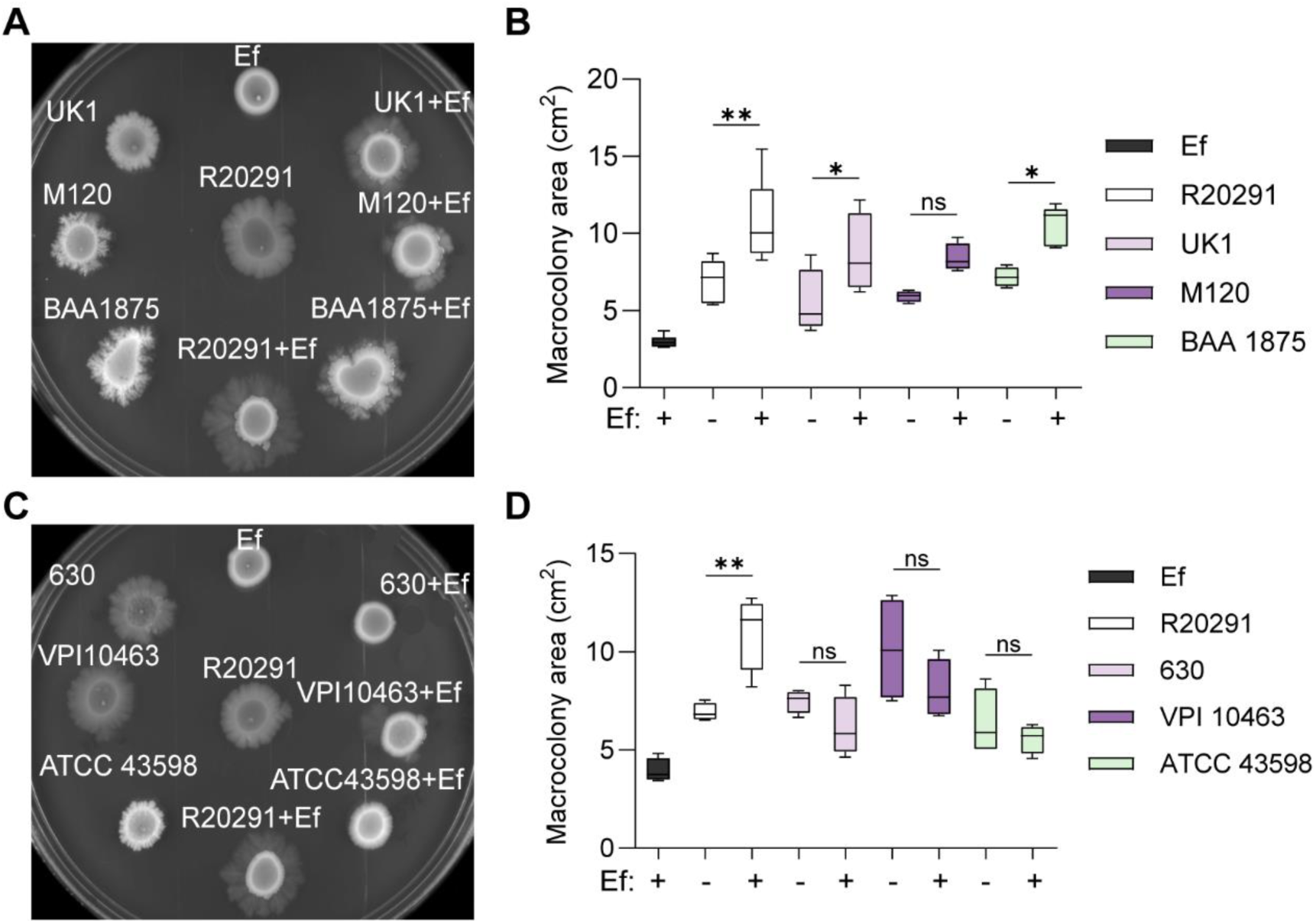
*E.* faecalis-driven increases in *cmr-*ON and *cmr-*phenotypes are not ubiquitous in *C. difficile* strains. (**A**) Representative image of tendril formation after 5 days of growth. *C. difficile* strains R20291 (RT027), UK1 (RT027), M120 (RT078), and BAA1875 (RT078) in monocultures and dual cultures with *E. faecalis* OG1RF at a 1:1 ratio. (**B**) Macrocolony area from (A) on day 5 of surface growth. (**C**) Representative image of tendril formation after 5 days of growth. *C. difficile* strains R20291 (RT027), 630 (RT012), VPI 10463 (RT087), and ATCC 43598 (RT017) in monocultures and dual cultures with *E. faecalis* OG1RF at a 1:1 ratio. (**D**) Macrocolony area from (D) on day 5 of surface growth. Data from independent experiments is presented as means and standard error. ** p < 0.01, * p < 0.05, one-way ANOVA with Šídák’s multiple comparison test.

Enteric infections are often polymicrobial by nature as the large intestine is inhabited by a dynamic and complex microbial community (21). Our previous work demonstrated that enterococcal-mediated changes in *C. difficile* virulence were specific to the enterococci and not conserved across all enteric bacteria (11). To test if the increased inversion towards *cmr-*ON in *C. difficile* R20291 stimulated by *E. faecalis* OG1RF was strain or species-specific, we first evaluated the ability of other enterococci to stimulate surface motility of *C. difficile* R20291 and to enrich for the *cmr-*ON state during co-culture on agar surfaces. While colony morphology was variable, we observed a significant shift to cmr-ON (60% to 95%) in *C. difficile* R20291 when cultured with *E. faecalis* strains OG1RF, an *E. faecalis* isolate from a microbial consortium generated from neonatal mice termed Pedscom, and the V583 vancomycin-resistant *E. faecalis* strain, compared to *C. difficile* in monoculture (**Figure 6A-B**) (22). We also evaluated the effect of non-*E. faecalis* enterococci in the shift towards *cmr-*ON in *C. difficile* R20291. A significant increase in *cmr-*ON was not achieved when *C. difficile* R20291 was cultured with multiple strains of *Enterococcus faecium* (Efm VRE and Efm DVT 705), *Enterococcus durans* (Ed), or endogenous *Enterococcus* selectively isolated from mice (murine Ent).

**Figure 6.**
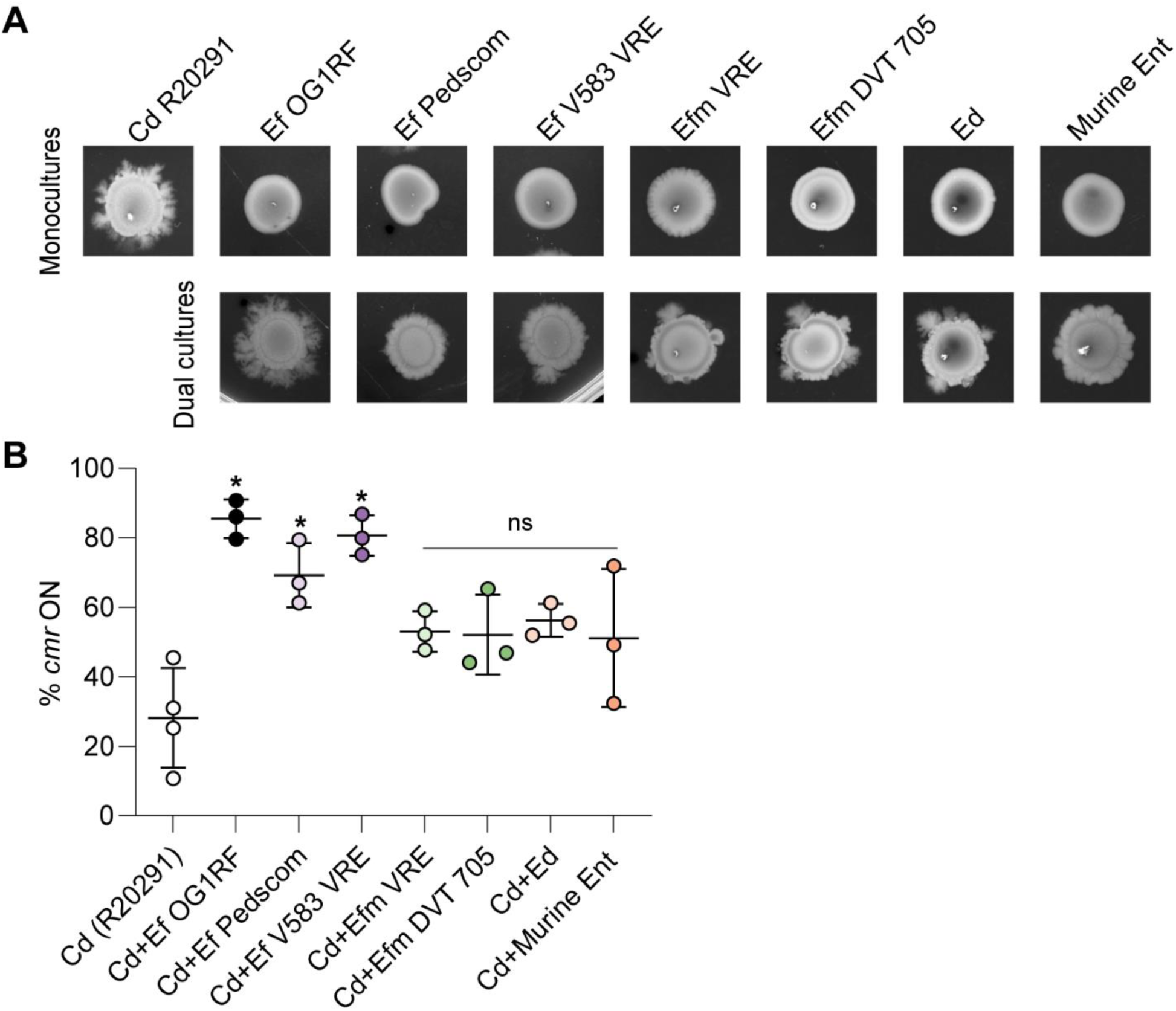
*E. faecalis* strains increase *cmr-*ON and *cmr-*phenotypes in *C. difficile.* (**A**) Representative image of surface motility after 5 days of growth. Surface motility of Cd R20291, Ef OG1RF, Ef Pedscom, Ef V583 VRE, Efm VRE, Efm DVT 705, Ed, and Murine Ent in monocultures (top) or dual cultures (bottom) with Cd R20291 at a 1:1 ratio. (**B**) Quantitative PCR of *cmr* switch orientation in macrocolonies from panel A after 5 days of growth with comparisons made to Cd. Data are presented as means and standard error, with symbols representing biological replicates. *** p < 0.001, Brown-Forsythe and Welch ANOVA tests (B, D).

Additionally, we were interested in whether bacteria outside of the *Enterococcus* genus were capable of enriching for *cmr*-ON in *C. difficile* R20291. To investigate this, we tested a consortium of enteric bacteria including *Staphylococcus epidermidis*, *Streptococcus agalactiae*, *Escherichia coli* MG1655, *E. coli* CE23, and *Enterobacter cloacae*. While *S. epidermidis* and *S. agalactiae* did not enrich for *cmr-*ON *C. difficile*, *E. coli* MG1655 and *E. coli* CE23 did lead to *cmr*-ON enrichment (**Supplemental Figure 3**). *E. cloacae* also appeared to enrich for *cmr*-ON, though the effect was not statistically significant. Further studies are required to better understand the extent to which enteric microbes, especially the Enterobacteriaceae, modulate phenotypic heterogeneity and phase variable systems in *C. difficile.* Although these experiments are limited to the bacterial strains tested, these observations point towards a preferential interaction of *E. faecalis* strains and select enteric bacteria with certain *C. difficile* strains that stimulates a shift to the *cmr-*ON state in *C. difficile*.

## DISCUSSION

Phase variation entails rapid phenotypic changes in bacterial populations as they face internal and external pressures, which are prevalent in dynamic environments like the large intestine. In this study, we investigated modulation of the phase variable signal transduction system CmrRST in the hypervirulent strain *C. difficile* R20291 by the enterococcal species *E. faecalis* OG1RF. We determined that *E. faecalis* can enrich the *cmr*-ON *C. difficile* population, altering broad phenotypes in *C. difficile*, including motility and cell morphology.

This investigation revealed enhanced tendril formation and surface motility in *C. difficile* R20291 when grown on solid agar in proximity to or in dual culture with *E. faecalis*. We found that this response leads to a change in phenotypes that is dependent on the orientation of the *cmr* switch. While *cmrR* and *cmrS* were not necessary for the phenotypic shift in *C. difficile*, *cmrT* was required for the change, consistent with previous work showing that while *cmrR* contributes to surface motility, *cmrT* is crucial (7, 9). Since the regulation of CmrRST is complex, we confirmed that *E. faecalis* does not detectably alter global levels of c-di-GMP in *C. difficile* to impact expression of *cmrRST*. However, it remains possible that localized fluxes of c-di-GMP in *C. difficile* cells or a subset of cells may influence CmrRST phenotypes. Autoregulation of *cmrRST* by CmrR also does not play a role, as deletion of *cmrR* did not eradicate the phenotypic switch in *C. difficile* dual culture with *E. faecalis*. These observations provide evidence of an inter-species interaction resulting in phase variation in *C. difficile* to the *cmr*-ON state.

Our previous and current work demonstrates that the CmrT response regulator is required for *C. difficile* spreading on solid surfaces, a phenotype associated with the *cmr-*ON population (7, 9). A selective advantage for *cmr*-ON *C. difficile* cells allows them to expand and become a greater proportion of the population. Here, we found that *cmrT* is necessary for shifting to *cmr*-ON in monoculture and in dual culture with *E. faecalis* on agar plates. The surface motility defect of Δ*cmrT* likely limits the opportunity to select for *cmr*-ON cells on solid surfaces. It remains possible that there is a more complex role for *cmrT* in favoring the *cmr-*ON orientation of the *cmr* switch independent of *E. faecalis* that requires further investigation.

A potential mechanism of selection for *cmr-*ON *C. difficile* is through *E. faecalis*-driven changes to the environment (24). Rough and smooth *C. difficile* colony morphotypes carry out distinct active metabolic reactions, and phase variation of CmrRST is sensitive to carbohydrate availability (24). Nutrient availability or deficiency can shift metabolic pathway utilization and could impact the fitness of the *cmr*-ON and *cmr*-OFF variants. Since *E. faecalis* has been shown to alter the metabolic environment in culture with *C. difficile,* these environmental shifts may drive selection for *cmr*-ON *C. difficile* phase variants (11). Determining the signal leading to *cmr*-ON enrichment is an area of future study for our labs.

We must also consider whether *E. faecalis* benefits from enrichment of *cmr*-ON populations of *C. difficile.* Since *cmr-*ON *C. difficile* has increased surface motility, it may be more advantageous in polymicrobial environments with *E. faecalis,* such as dual-species biofilms (11). Furthermore, rough colony isolates of *C. difficile* displayed heightened virulence compared to smooth isolates in a hamster model of infection, so it is possible that *E. faecalis* modulation of *cmr*-associated behaviors also contributes to the enhanced virulence observed in *C. difficile-E. faecalis* co-infections (7, 11). The effect of *E. faecalis* on CmrRST *in vivo* is a pertinent area for future study. While our work began to address this through assessing if one endogenous murine enterococcal strain can favor *cmr*-ON *C. difficile*, there is much left to be understood. This includes determining if *E. faecalis* impacts *cmr* switch orientation in *C. difficile* populations during CDI, and if so, how this impacts toxin-independent pathogenesis behaviors of *C. difficile.* Additionally, this presents an opportunity to examine the metabolic niches in which *C. difficile* and *E. faecalis* reside and how the ecological context in the GI tract impacts CDI outcomes. This is especially relevant to our findings that *E. faecium* does not elicit the same behaviors in *C. difficile* as *E. faecalis* and other enteric pathogens. Our previous work highlights how *E. faecalis* and *E. faecium* occupy varied metabolic niches and affect *C. difficile* behavior, including toxin production (11). These results emphasize inter-species interactions that require further investigation into their mechanisms and clinical relevance.

Our work underscores that bacterial behaviors are controlled by a complex network including microbes, surfaces, and the environment (6, 7, 11). In this case, these microbe-driven environmental factors alter *C. difficile* phenotypes, which can directly impact its fitness. Our study contributes to research investigating how phenotypically distinct populations and phase variation influence pathogen physiology and behaviors, and potentially infection. For example, one of the two alternating subpopulations formed by *Acinetobacter baumannii* is enriched in tissues and associated with worse outcomes during infection (25). In addition, recent metagenomic analysis has provided evidence of epigenetic phase variation within the intestinal microbiota (26). The modulation of *C. difficile* phase variation by *E. faecalis* may be central to virulence during CDI. Hence, microbial responses and environmental factors are increasingly important when considering *C. difficile* treatment and vaccination strategies, where existing microbiota members may impact *C. difficile* behavior, accessibility to the host, and virulence.

This work suggests that microbiota structure and ecological context in the gut are critical for controlling *C. difficile* phenotypic heterogeneity. Altogether, these results demonstrate that in the context of complex ecosystems, like those experienced by invading pathogens during enteric infections, bystander bacteria can influence phase variation to promote phenotypes that may be advantageous to survival, niche establishment, or pathogenesis. Our work establishes an important paradigm that microbial ecology and context can markedly reshape pathogen behavior through phase variation.

## MATERIALS AND METHODS

### Growth and maintenance of bacterial strains

Table S1 lists the strains and plasmids used in this study. *C. difficile* and *E. faecalis* strains were grown in an anaerobic environment of 90% N_2_, 5% CO_2_, and 5% H_2_. The strains were statically grown at 37°C in brain heart infusion supplemented with yeast extract (BHIS) + L-cysteine broth or BHIS agar (BHISA) + L-cysteine (37 g/L Bacto brain heart infusion or brain heart infusion agar, 5 g/L yeast extract).

### Generation of the *fliC* mutant

The *fliC* gene (CDR20291_0240) was deleted in-frame using an allelic exchange method described previously (27, 28). Regions of homology upstream and downstream of fliC were amplified from R20291 genomic DNA using primers fliC-F1 + fliC-R1 and fliC-F2 + fliC-R2, respectively. The fragments were cloned into pMSR0 at the BamHI site using a Gibson assembly (New England Biolabs). Reaction products were transformed into *Escherichia coli* DH5α and plated on Luria-Bertani (LB)-agar containing 20 µg/mL chloramphenicol (Cm). Cm-resistant colonies were screened by PCR using primers fliC-F1 and fliC-R2 and confirmed by sequencing. The resulting pMSR0::ΔfliC plasmid was introduced into *C. difficile* R20291 via conjugation with *E. coli* HB101(pRK24), with plating on BHIS-agar containing 20 µg/mL thiamphenicol (Tm) and 100 µg/mL kanamycin (Kan). Tm-Kan-resistant transconjugants were subcultured as described (27, 28). Colonies were screened for the *fliC* deletion by PCR with primers fliC-F0 + fliC-R0 and were confirmed to be non-motile in BHIS-0.3% agar medium (29).

### Proximity growth assays on solid media

BHISA + L-cysteine plates were marked for inoculation with two dots 0.5 cm apart, and additional dots were added at 0.5cm apart. Overnight cultures prepared in BHIS were normalized to an OD of 0.5. For the inoculum, 10 µL of the normalized cultures of *E. faecalis* and *C. difficile* strains were spotted on opposite sides. Plates grew for 7 days before imaging.

### Surface motility assays

The optical density (OD) of overnight cultures (16 - 20 hrs) prepared in BHIS broth was measured through absorbance (600 nm). The cultures were normalized with fresh BHIS to an OD_600_ of 0.5, and dual cultures were prepared at 1:1. From the monocultures and dual cultures, 10 µL was spotted onto BHISA + L-cysteine. Plates were imaged in Syngene G: Box Imager or ChemiDoc Imager, and areas of the macrocolonies were quantified using ImageJ (version 1.54j).

### Quantitative reverse transcriptase PCR

For gene expression analysis in *C. difficile*, colonies were collected, suspended in 1 mL of acetone: ethanol, and stored at - 80°C prior to RNA isolation. RNA was extracted as described in previous work (30) and treated with the TURBO DNA-*free* Kit (Invitrogen^TM^) to remove contaminating genomic DNA. cDNA synthesis was performed using the high-capacity cDNA Reverse Transcription Kit (Applied Biosystems^TM^) according to the manufacturer’s protocol. The quantitative PCR was performed using SensiFAST SYBR and fluorescein kit (Bioline) as previously described (7, 8). Data were analyzed using *rpoC* as a reference gene and normalization to the indicated strain. The primers used are defined in Table S2.

### Quantification of *cmr* switch orientation by qPCR

Samples were collected, and genomic DNA was extracted and purified by phenol:chloroform:isopropanol and ethanol precipitation. qPCR was performed as previously described with 100 ng of DNA per 20 μL reaction and 100 nM primers (7). Quantification was performed using the ΔΔ*CT* method, with *rpoC* as the reference gene. The primers used are defined in Table S2.

### Imaging rough and smooth colonies

To image colony morphology from *C. difficile* cultured with or without *E. faecalis*, surface motility assays were performed as described above. After incubation, macrocolonies were collected and resuspended in 0.5 mL of 1X DPBS. Serial dilutions (1:10) were plated on BHIS + L-cysteine plates and incubated for 48 hours. Images of plates were taken at 2X with the Syngene G: Box Imager.

### Microscopy

Scanning electron microscope (SEM) experiments were carried out at CDB Microscopy Core (Dept. of Cell and Developmental Biology at the Perelman School of Medicine, University of Pennsylvania). Bacterial macrocolonies were grown on BHISA for 7 days, excised from the petri dish, and inserted into a 24-well plate with a 12 mm standard coverslip underneath the agar. Samples were washed with PBS and fixed in 50 mM Na-cacodylate buffer overnight. Samples were washed three times with 50 mM Na-cacodylate buffer, fixed for 2 hours with 2.5% glutaraldehyde in 50 mM Na-cacodylate buffer (pH 7.3), and then dehydrated in a graded series of ethanol concentrations through 100% for 1.5 hours. Dehydration in 100% ethanol was done three times. After dehydration, samples were incubated for 20 minutes in 50% HMDS in ethanol, followed by three changes of 100% HMDS (Sigma-Aldrich Co.), followed by overnight air-drying as described previously (31). Samples were mounted on stubs and sputter-coated with gold palladium. Specimens were observed and imaged using a Quanta 250 FEG scanning electron microscope (FEI, Hillsboro, OR, USA) at 10 kV accelerating voltage.

Gram stains were performed on the edges of macrocolonies collected from the surface motility assays. Stained cells were visualized using a Keyence BZ-X810 microscope. For quantification of cell length, images from biological replicates were analyzed using ImageJ.

Fluorescence microscopy was performed as previously described (20). Surface motility assays were performed as described above with *C. difficile* c-di-GMP reporter strains p*_gluD-_*PRS::mCherryOpt and p*_gluD_*_-_PRS^A70G^::mCherryOpt (negative control) in wildtype backgrounds. Macrocolonies of mono- and dual cultures were collected and resuspended in 1 mL of 1X DPBS, pelleted, and resuspended in 0.5 mL of 1X DPBS. Samples were combined with 120 µL of fixative (100 µL 16% PFA and 20 µL 1M NaPO_4_ pH 7.4) and incubated for 30 minutes at room temperature covered from light. Samples were applied to 1% agarose pads for microscopy using a Keyence BZ-X810 equipped with Chroma 49005-UF1 for RFP detection and a 100x oil immersion Nikon Plan Apo objective.

### Human samples

For the DYNAMIC study, subjects were recruited at the Children’s Hospital of Philadelphia (CHOP) from September 2015 to April 2018 and informed consent was acquired (IRB approval number 15-011817), as previously described (11, 32).

### Reproducibility and statistical analysis

By conducting and cross-validating experiments at two independent institutions, we confirmed findings were robust and replicable across conditions and environments. Statistical analyses were performed using GraphPad Prism version 10.4.2. Specific statistical tests, replicate numbers, calculated errors, and extended information for each experiment are reported in the figure legends. All data represent distinct samples unless otherwise stated.

## ACKNOWLEDGMENTS

We thank the members of the Zackular and Tamayo laboratories for their support and critical feedback on this manuscript. We thank Daria Van Tyne, Paul Planet, Mark Goulian, Aaron Hecht, and Kelly Doran for sharing bacterial strains used in this publication. We thank Eric Skaar and the Skaar laboratory for helpful discussions regarding this work. Thank you to the CDB Microscopy Core at the University of Pennsylvania for their assistance with SEM imaging.

This work was supported by NIH awards R01-AI143638 and R01-AI188648 to R.T. and GRT-00000565, GRT-00003300-0226, and K22AI7220 to J.P.Z. A.S.W. was supported by the NSF GRFP. The funders had no role in study design, data collection and analysis, decision to publish, or preparation of the manuscript.

---

**Supplemental Figure 1.**
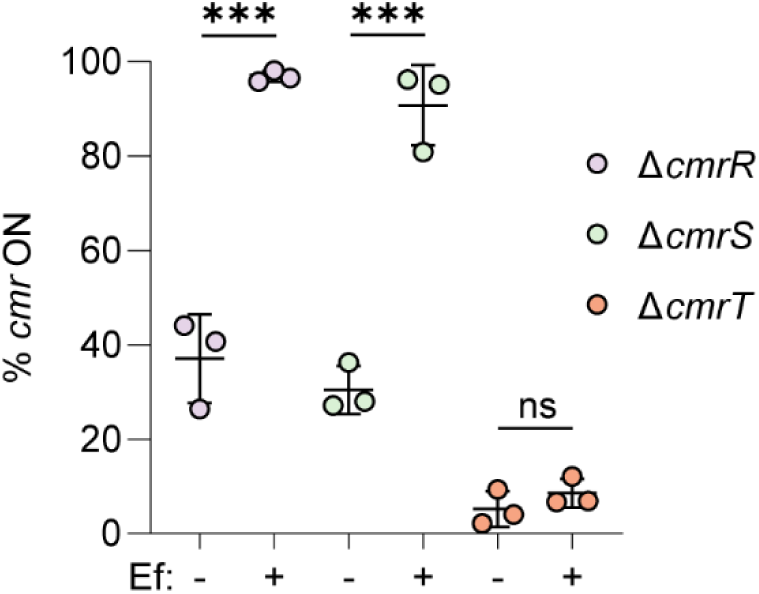
*C. difficile* requires *cmrT* for the *E. faecalis*-mediated increase in *cmr*-ON. Quantitative PCR of *cmr* switch orientation in macrocolonies at after 5 days of growth of Δ*cmrR* Cd, Δ*cmrS* Cd, and Δ*cmrT* Cd in monocultures or dual cultures with Ef at a 1:1 ratio. Data from 3 biological replicates were analyzed and presented as means and standard error. *** p < 0.001, unpaired t-tests.

**Supplemental Figure 2.**
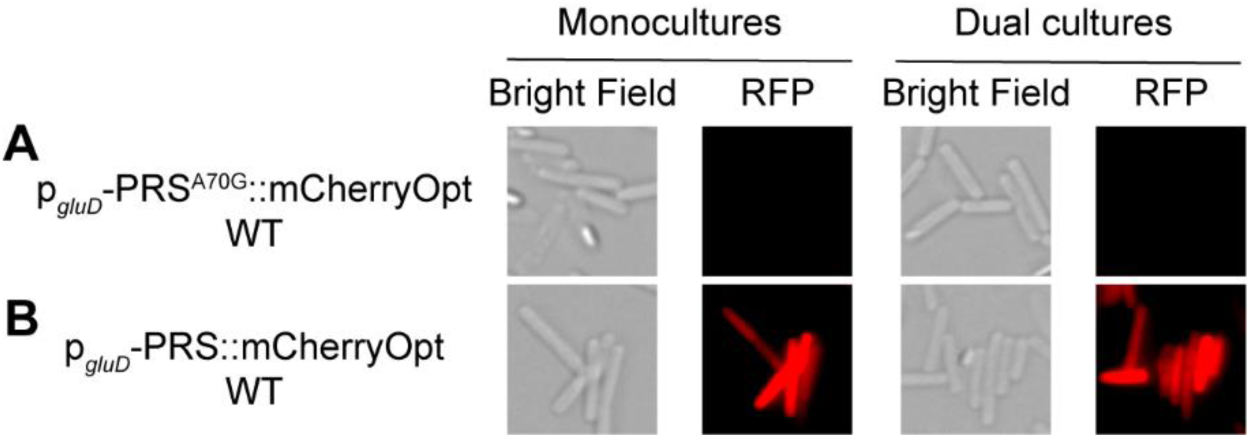
*E. faecalis* does not alter global c-di-GMP levels in culture with *C. difficile*. Representative micrographs of reporter strains. *C. difficile* reporter strains (**A**) p*_gluD_*_-_PRS^A70G^::mCherryOpt (negative control), and (**B**) p*_gluD-_* PRS::mCherryOpt, were grown for 7 days in monocultures or dual cultures with *E. faecalis* at a 1:1 ratio before preparing micrographs and imaging at 100X.

**Supplemental Figure 3.**
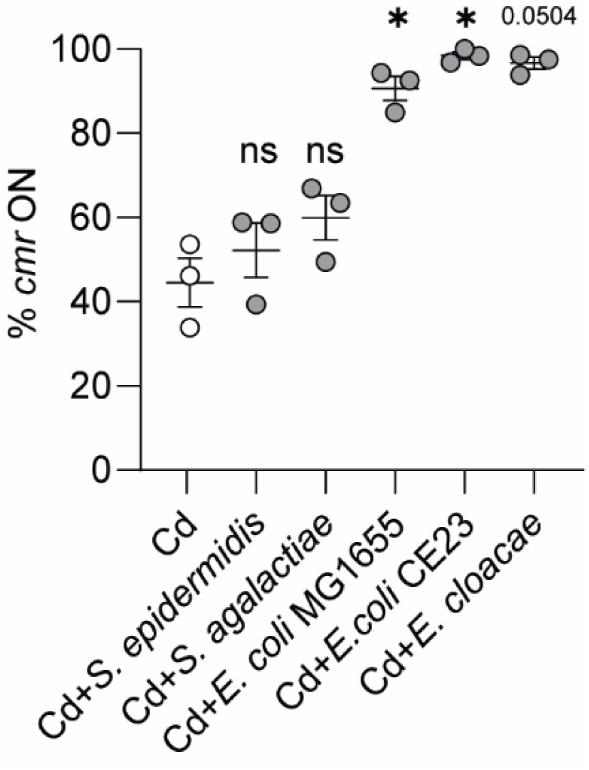
Certain enteric bacterium increase *cmr*-ON in *C. difficile.* Quantitative PCR of *cmr* switch orientation in macrocolonies after 5 days of growth with comparisons made to Cd. Macrocolonies were Cd monocultures or dual cultures with Cd and *S. epidermidis*, *S. agalactiae*, *E. coli* MG1655, *E. coli* CE23, or *E. cloacae* at a 1:1 ratio. Data are presented as means and standard error, with symbols representing biological replicates. * p < 0.05, Brown-Forsythe and Welch ANOVA tests.

**Table S1.** Strains and plasmids used in this study.

| Lab<br>Notation* | Strain name | Description | Reference |
| --- | --- | --- | --- |
| ***C. difficile* strains** |  |  |  |
| RT273,<br>Z033 | *C. difficile* R20291 | Ribotype 027 strain, smooth colony isolate (BioSample GCA_000027105) | (33) |
| RT3332 | *C. difficile* $\Delta fliC$ | R20291 with in-frame deletion of *fliC* | This work |
| RT1341 | *C. difficile* $\Delta pilA$ | R20291 *pilA1::ermB*; Targetron Insertion | (34) |
| RT1566 | *C. difficile* $\Delta sigD$ | R20291 *sigD::ermB*; Targetron Insertion | (35) |
| RT947 | *C. difficile* $\Delta pilB$ | R20291 *pilB1::ermB*; Targetron Insertion | (16) |
| RT2256,<br>Z036 | *C. difficile* $\Delta cmrR$ | R20291 with in-frame deletion of *cmrR* | (7) |
| RT2807,<br>Z209 | *C. difficile* $\Delta cmrS$ | R20291 with in-frame deletion *cmrS* | (7) |
| RT2257,<br>Z037 | *C. difficile* $\Delta cmrT$ | R20291 with in-frame deletion of *cmrT* | (7) |
| RT2296,<br>Z038 | *C. difficile* $\Delta cmrR \Delta cmrT$ | R20291 with in-frame deletions of *cmrR* and *cmrT* | (7) |
| RT2406,<br>Z210 | *C. difficile* *cmr*Δ3-ON | R20291 with *cmr* switch phase-locked in ON orientation due to deletion of three nucleotides in the right inverted repeat | (9) |
| RT2395,<br>Z211 | *C. difficile* *cmr*Δ3-OFF | R20291 with *cmr* switch phase-locked in OFF orientation due to deletion of three nucleotides in the right inverted repeat | (9) |
| RT2816 | *C. difficile*<br>*P<sub>gluD</sub>*·PRS::mCherryOpt | R20291 with c-di-GMP reporter consisting of *gluD* promoter, *pilA1* c-di-GMP riboswitch, and codon-optimized mCherry | (20) |
| RT2813 | *C. difficile*<br>*P<sub>gluD</sub>*-<br>PRS<sup>A70G</sup>::mCherryOpt | R20291 with c-di-GMP reporter containing a mutation (A70G) in the riboswitch rendering it blind to c-di-GMP | (20) |
| RT1065 | *C. difficile* UK1 | Ribotype 027 | (36, 37) |
| RT3321 | *C. difficile* M120 | Ribotype 078 | (38) |
| RT1357 | *C. difficile* ATCC BAA-1875 | Ribotype 078 | ATCC |
| RT1124 | *C. difficile* 630 | Ribotype 012 strain | (7) |
| RT1125 | *C. difficile* VPI 10463 | Ribotype 087 strain | (39) |
| RT1358 | *C. difficile* ATCC 43598 | Ribotype 017 strain | ATCC |
| ***Enterococcus* strains** |  |  |  |
| RT2378,<br>Z002 | *E. faecalis* OG1RF | “EF OG1RF”; ATCC 47077 | ATCC |
| Z007 | *E. faecalis* V583 VRE | “Ef V583 VRE”; ATCC 700802 | ATCC |
| ABS055 | *E. faecalis* | “Ef Pedscom”; strain from Pedscom | (22) |
|  | Pedscom PC15 | defined microbial consortia shared by CHOP |  |
| ABS078 | *E. faecium* VRE | "Efm VRE". ATCC 700221 | ATCC |
| Z221 | *E. faecium* DVT705 VRE | "Efm DVT705"; clinical isolate from the University of Pittsburgh | (40) |
| AW25 | *E. durans* | "Ed"; isolated from clinical sample in DYNAMIC study, DYNAMIC 1-16 | (32) |
| AW40 | Murine *Enterococcus* | "Murine Ent"; *Enterococcus* isolated in the Zackular lab from cefoperazone and vancomycin-treated mouse stool using bile esculin plates | This work |
| **Enteric bacterial strains** |  |  |  |
| Z094,<br>AW50 | *E. coli* MG1655 | Substrain of *E. coli* K-12 from UPenn | NCBI: txid511145 |
| Z178,<br>AW54 | *E. coli* CE23 | Enteropathogenic *E. coli* clinical strain from UPenn |  |
| AW43 | *S. agalactiae* CJB111 | "*S. agalactiae*"; strain from the University of Colorado Anschutz | NCBI: txid342617 |
| Z227,<br>AW38 | *S. epidermidis* | ATCC 14490 | ATCC |
| Z074 | *E. cloacae* | ATCC 13047 | ATCC |

\* RT notation indicates strain in R. Tamayo lab collection; Z/AW notation indicates strain in J. Zackular lab collection.

**Table S2.**
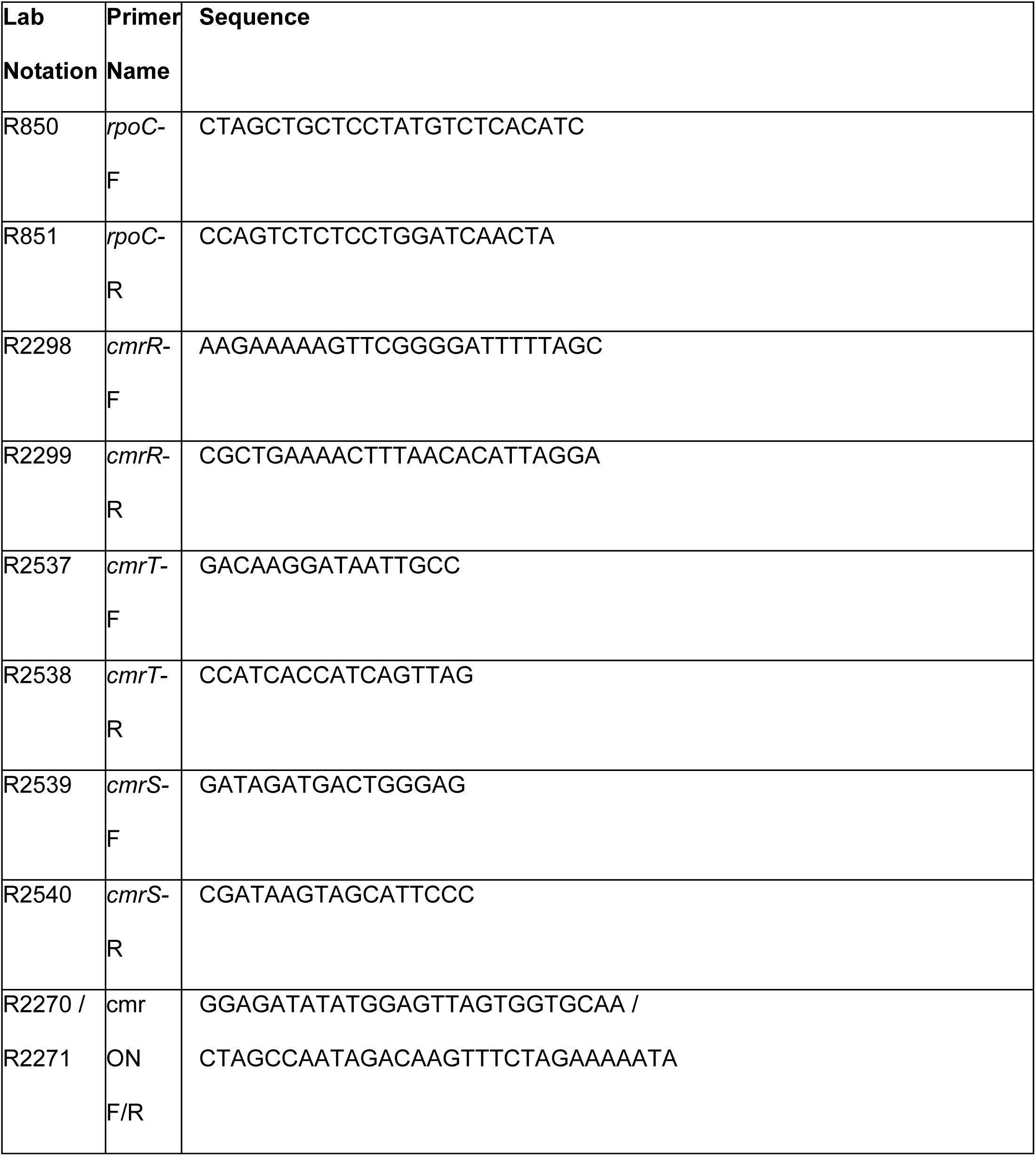
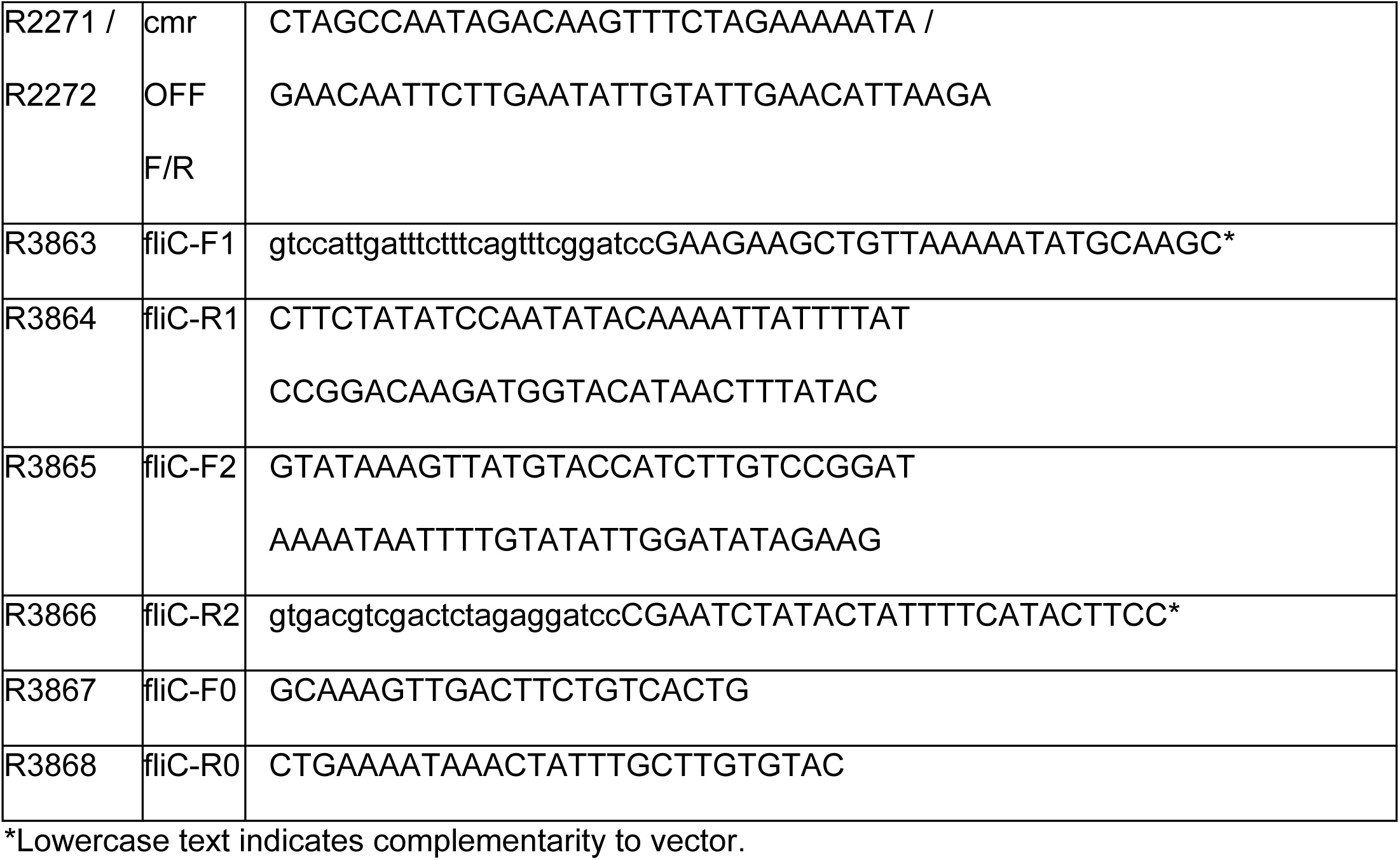
Primers used in this study.

